# A method for identifying genetic heterogeneity within phenotypically-defined disease subgroups

**DOI:** 10.1101/037713

**Authors:** James Liley, John A Todd, Chris Wallace

## Abstract

Many common diseases show wide phenotypic variation. We present a statistical method for determining whether phenotypically defined subgroups of disease cases represent different genetic architectures, in which disease-associated variants have different effect sizes in the two subgroups. Our method models the genome-wide distributions of genetic association statistics with mixture Gaussians. We apply a global test without requiring explicit identification of disease-associated variants, thus maximising power in comparison to a standard variant by variant subgroup analysis. Where evidence for genetic subgrouping is found, we present methods for post-hoc identification of the contributing genetic variants.

We demonstrate the method on a range of simulated and test datasets where expected results are already known. We investigate subgroups of type 1 diabetes (T1D) cases defined by autoantibody positivity, establishing evidence for differential genetic architecture with thyroid peroxidase antibody positivity, driven generally by variants in known T1D associated regions.

## Introduction

Analysis of genetic data in human disease typically uses a binary disease model of cases and controls. However, many common human diseases show extensive clinical and phenotypic diversity which may represent multiple causative pathophysiological processes. Because therapeutic approaches often target disease-causative pathways, understanding this phenotypic complexity is valuable for further development of treatment, and the progression towards personalised medicine. Indeed, identification of patient subgroups characterised by different clinical features can aid directed therapy [1] and accounting for phenotypic substructures can improve ability to detect causative variants by refining phenotypes into subgroups in which causative variants have larger effect sizes [2].

Such subgroups may arise from environmental effects, reflect population variation in non-disease related anatomy or physiology, correspond to partitions of the population in which disease heritability differs, or represent different causative pathological processes. Our method tests whether there exist a subset of disease-associated SNPs which have different effect sizes in case subgroups, determining whether heterogeneity corresponds to differential genetic pathology.

Our test is for a stronger assertion than the question of whether subgroups of a disease group exhibit any genetic differences at all, as these may be entirely disease-independent: for example, although there will be systematic genetic differences between Asian and European patient cohorts with type 1 diabetes (T1D), these differences will not generally relate to the pathogenesis of disease.

Rather than attempting to analyse SNPs individually for differences between subgroups, a task for which GWAS are typically underpowered, we model allelic differences across all SNPs using mixture multivariate normal models. This can give insight into the structure of the genetic basis for disease. Given evidence that there exists some subset of SNPs that both differentiate controls and cases and differentiate subgroups, we can then reassess test statistics to search for single-SNP effects.

## Results

### Summary of proposed method

We jointly consider allelic differences between the combined case group and controls, and allelic differences between case subgroups independent of controls. Specifically, we establish whether the data support a hypothesis (*H*_1_) that a subset of SNPs associated with case-control status have different underlying effect sizes (and hence underlying allele frequencies) in case subgroups. This assumption has been used previously for genetic discovery [3].

*H*_1_ encompasses several potential underlying mechanisms of heterogeneity. A set of SNPs may be associated with one case subgroup but not the other; the same set of SNPs may have different relative effect sizes in subgroups, or heritability may differ between subgroups. These scenarios are discussed in supplementary note 1.

Our overall protocol is to fit two bivariate Gaussian mixture models, corresponding to null and alternative hypotheses, to summary statistics (*Z* scores) derived from SNP data. We assume a group of controls and two non-intersecting case subgroups, and jointly consider allelic differences between the combined case group and controls, and allelic differences between case subgroups independent of controls (figure 1). Heterogeneity in cases can also be characterised by a quantitative trait, rather than explicit subgroups.

**Figure 1:**
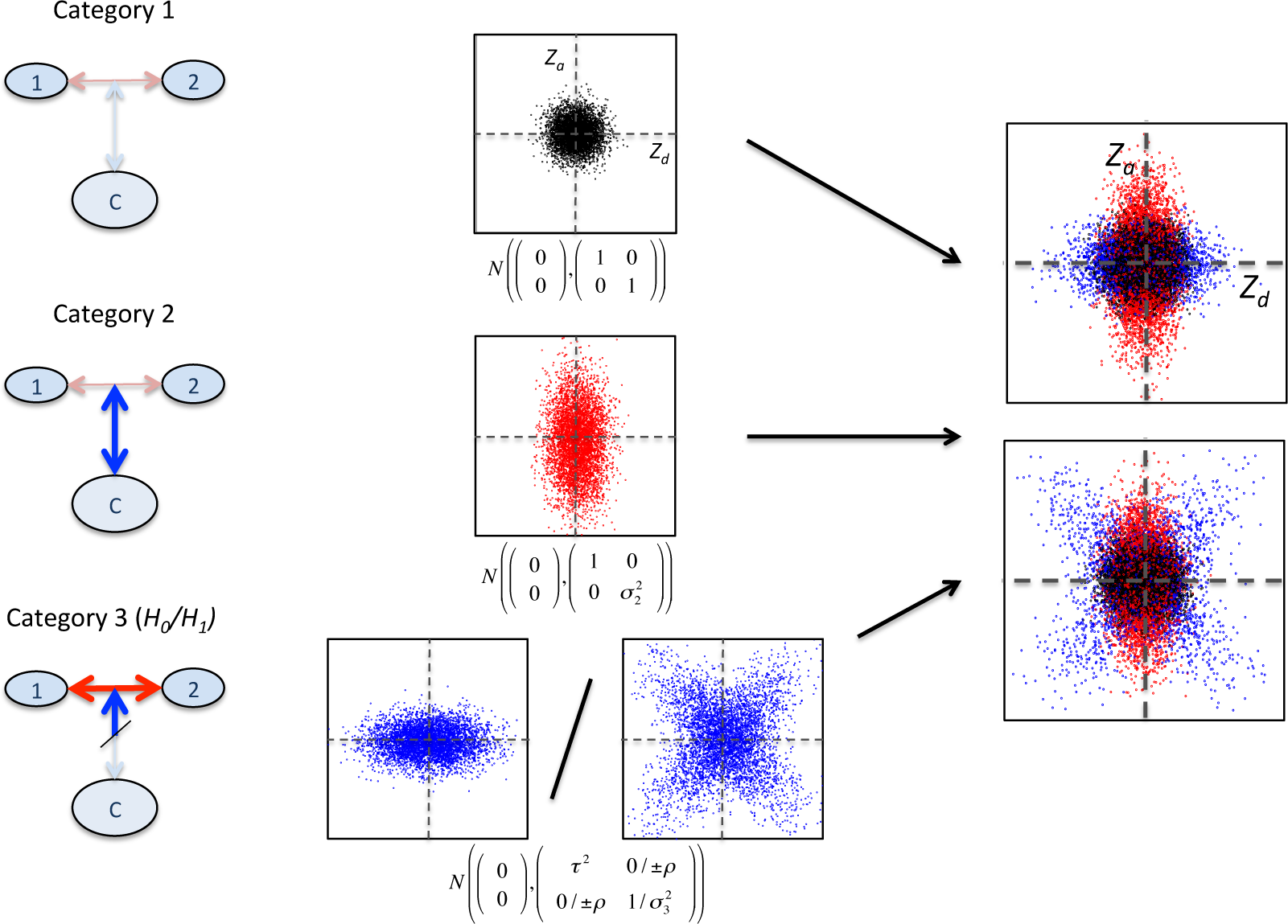
Overview of three-categories model. *Z*_*d*_ and *Z*_*a*_ are *Z* scores derived from GWAS p-values for allelic differences between case subgroups (1 vs 2), and between cases and controls (1+2 vs *C*) respectively (left). Within each category of SNPs, the joint distribution of (*Z*_*d*_, *Z*_*a*_) has a different characteristic form. In category 1, *Z* scores have a unit normal distribution; in category 2, the marginal variance of *Z*_*a*_ can vary. The distribution of SNPs in category 3 depends on the main hypothesis. Under *H*_0_ (that all disease-associated SNPs have the same effect size in both subgroups), only the marginal variance of *Z*_*d*_ may vary; under *H*_1_ (that subgroups correspond to differential effect sizes for disease-associated SNPs), any covariance matrix is allowed. The overall SNP distribution is then a mixture of Gaussians resembling one of the rightmost panels, but with SNP category membership unobserved. Visually, our test determines whether the observed overall *Z*_*d*_, *Z*_*a*_ distribution more closely resembles the bottom rightmost panel than the top.

For a given SNP we denote by *μ*_1_, *μ*_2_, *μ*_12_ and *μ*_*c*_ the population minor allele frequencies for each of the two case subgroups, the whole case group and the control group respectively, and *P*_*d*_, *P*_*a*_ GWAS p-values for comparisons of allelic frequency between case subgroups and between cases and controls, under the null hypotheses *μ*_1_ = *μ*_2_ and *μ*_12_ = *μ*_*c*_ respectively (or similarly for quantitative heterogeneity). We then derive absolute *Z* scores |*Z_d_*| and |*Z_a_*| from these p-values (see figure 1). We consider the values |*Z_d_*|, |*Z_a_*| as absolute values of observations of random variables (*Z*_*d*_, *Z*_*a*_) which are samples from a mixture of three bivariate Gaussians. Further details are given in supplementary note 2.

We consider each SNP to fall into one of three categories, with each category corresponding to a different joint distribution of *Z*_*d*_, *Z*_*a*_:

1. SNPs which do not differentiate subgroups and are not associated with the phenotype as a whole (*μ*_*c*_ = *μ*_*1*_ = *μ*_*2*_)
2. SNPs which are associated with the phenotype as a whole but which are not differentially associated with the subgroups (*μ*_*c*_ ≠ *μ*_12_; *μ*_1_ = *μ*_2_ = *μ*_12_)
3. SNPs which have different population allele frequencies in subgroups, and may or may not be associated with the phenotype as a whole (*μ*_1_ ≠ *μ*_2_)

If the SNPs in category 3 are not associated with the disease as a whole (null hypothesis, *H*_0_), we expect *Z*_*d*_, *Z*_*a*_ to be independent and the variance of *Z*_*a*_ to be 1. If SNPs in category 3 are also associated with the disease as a whole (alternative hypothesis, *H*_1_), the joint distribution of (*Z*_*d*_, *Z*_*a*_) will have both marginal variances greater than 1, and *Z*_*a*_, *Z*_*d*_ may co-vary. Our test is therefore focussed on the form of the joint distribution of (*Z*_*d*_, *Z*_*a*_) in category 3. Importantly, we allow that the correlation between *Z*_*d*_ and *Z*_*a*_ may be simultaneously positive at some SNPs and negative at others. This allows for a subset of SNPs to specifically alter risk of one subgroup, and another subset to alter risk for the other subgroup. To accommodate this, we only consider absolute *Z* scores and model the distribution of SNPs in category 3 with two mirror-image bivariate Gaussians.

Amongst SNPs with the same frequency in disease subgroups (categories 1 and 2), *Z*_*a*_ and *Z*_*d*_ are independent and the expected standard deviation of *Z*_*d*_ is 1. We therefore model the overall joint distribution of (*Z*_*d*_, *Z*_*a*_) as a Gaussian mixture in which the *pdf* of each observation (*Z*_*d*_, *Z*_*a*_) is given by

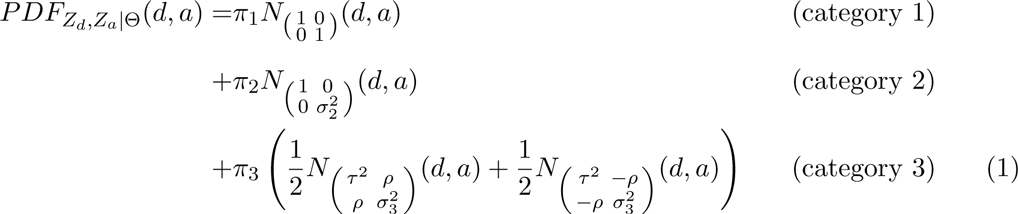

where *N*_∑_(*d*, *a*) denotes the density of the bivariate normal *pdf* centered at 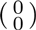 with covariance matrix ∑ at (*d*, *a*). Θ is the vector of values (*π*_1_, *π*_2_, *τ*, *σ*_2_, *σ*_3_, *ρ*). Under *H*_0_, we have *ρ* = 0 and *σ*_3_ = 1. The values (*π*_1_, *π*_2_, *π*_3_) represent the proportion of SNPs in each category, with ∑*π_i_* = 1 (see table 1). Patterns of (*Z*_*d*_, *Z*_*a*_) for different parameter values are shown in supplementary table 1.

**Table 1:**
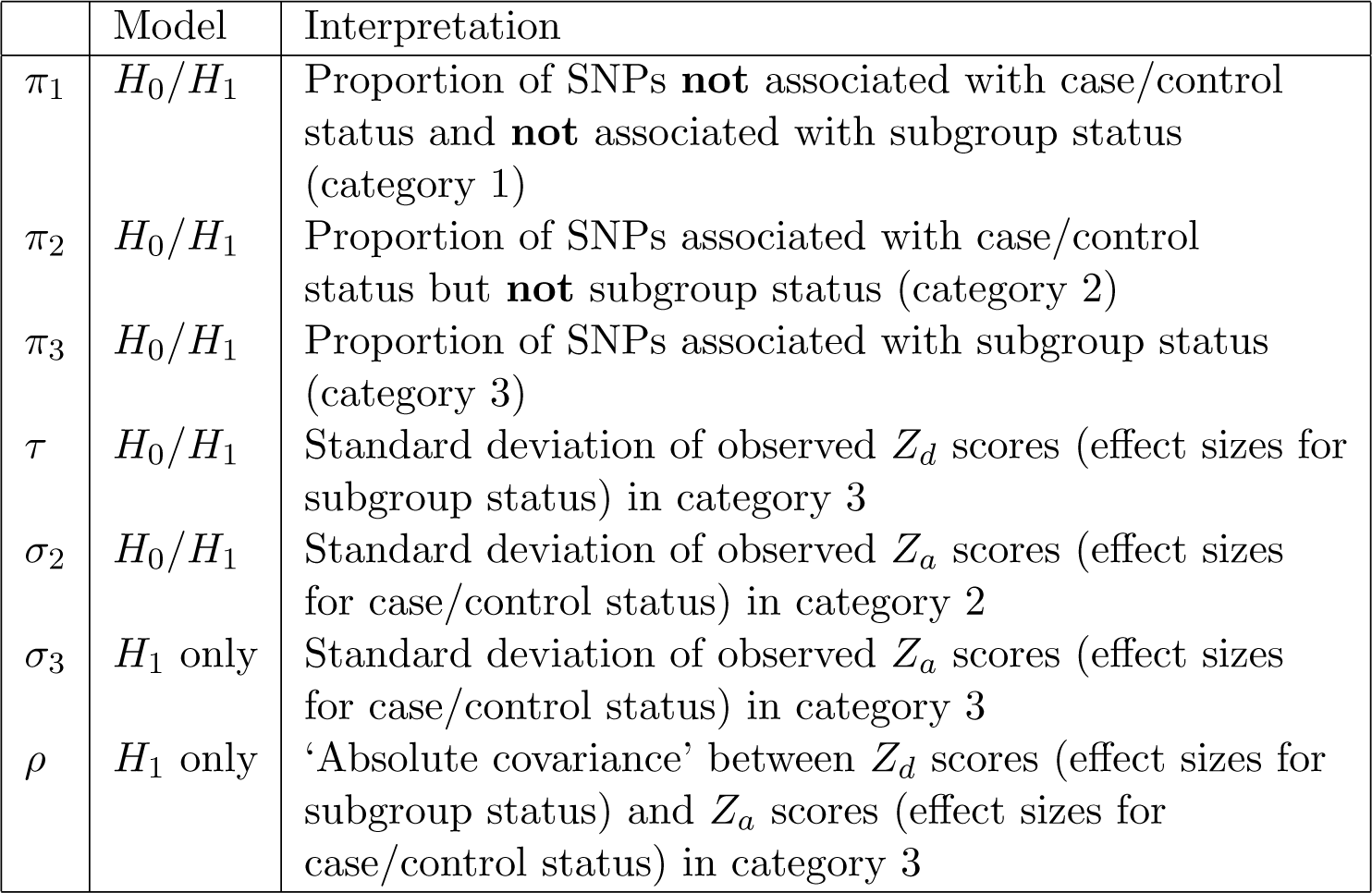
Interpretation of parameter values in the fitted model. Parameters *τ*, *σ*_2_ and *σ*_3_ are dependent on sample sizes, but can be converted to sample-size independent forms (see supplementary note, section 3.3)

We use the product of values of the above *pdf* for a set of observed *Z*_*d*_, *Z*_*a*_ as an objective function (‘pseudo-likelihood’, PL) to estimate the values of parameters. This is not a true likelihood as observations are dependent due to linkage disequilibrium (LD), although because we minimise the degree of LD between SNPs using the LDAK method [4], the PL is similar to a true likelihood.

### Model fitting and significance testing

We fit parameters *π*_1_, *π*_2_, *π*_3_ (= 1 − *π*_1_ − *π*_2_), *σ*_2_, *σ*_3_, *τ* and *ρ* under *H*_1_ and *H*_0_. Under *H*_0_, (*ρ*, *σ*_3_) = (0, 1).

We then compare the fit of the two models using the log-ratio of PLs, giving an unadjusted pseudo-likelihood ratio (uPLR). We subtract a term depending only on *Z*_*a*_ to minimise the influence of the *Z*_*a*_ score distribution, and add a term *log*(*π*_1_*π*_2_*π*_3_) to ensure the model is identifiable [5]. We term the resultant test statistic the pseudo-likelihood ratio (PLR). The distribution of the PLR is minorised by a distribution of the form:

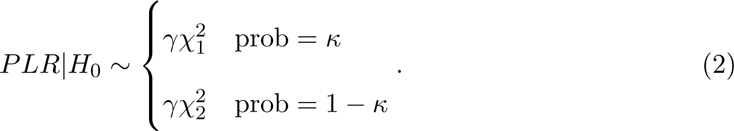

The value *γ* arises from the weighting derived from the LDAK procedure causing a scale change in the observed *PLR*. The mixing parameter *κ* corresponds to the probability that *ρ* = 0, (approximately 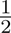).

We estimate *γ* and *κ* by sampling random subgroups of the case group. Such subgroups only cover the subspace of *H*_0_ with *τ* = 1 (no systematic allelic differences between subgroups), causing the asymptotic approximation of PLR by equation 2 to be poor. We thus estimate *γ* and *κ* from the distribution of a similar alternative test statistic, the cPLR (see methods section and supplementary note, section 2.5.1), which is well-behaved even when *τ* ≈ 1 and which majorises the distribution of PLR.

A natural next step is to search for the specific variants contributing to the PLR. An effective test statistic for testing subgroup differentiation for single SNPs is the Bayesian conditional false discovery rate (cFDR) [6, 7] applied to *Z_d_* scores ‘conditioned’ on *Z_a_* scores. However, this statistic alone cannot capture all the means by which the joint distribution of (*Z_a_, Z_d_*) can deviate from *H*_0_, and we also propose three other test statistics, each with different advantages, and compare their performance (supplementary note, section 5.1).

### Power calculations, simulations, and validation of method

We tested our method by application to a range of datasets, using simulated and resampled GWAS data. First, to confirm appropriate control of type 1 error rates across *H*_0_, we simulated genotypes of case and control groups under *H*_0_ for a set of 5×10^5^ autosomal SNPs in linkage equilibrium (supplementary note 3). Quantiles of the empirical PLR distribution were smaller than those for the empirical cPLR distribution and the asymptotic mixture-*χ*^2^, indicating that the test is conservative when *τ* > 1 (estimated type 1 error rate 0.048, 95% CI 0.039-0.059) and when *τ* ≈ 1 (estimated type 1 error rate 0.033, 95% CI 0.022-0.045) as expected; see figure 2. The distribution of cPLR closely approximated the asymptotic mixture-*χ*^2^ distribution across all values of *τ* (supplementary note, section 3.1).

**Figure 2:**
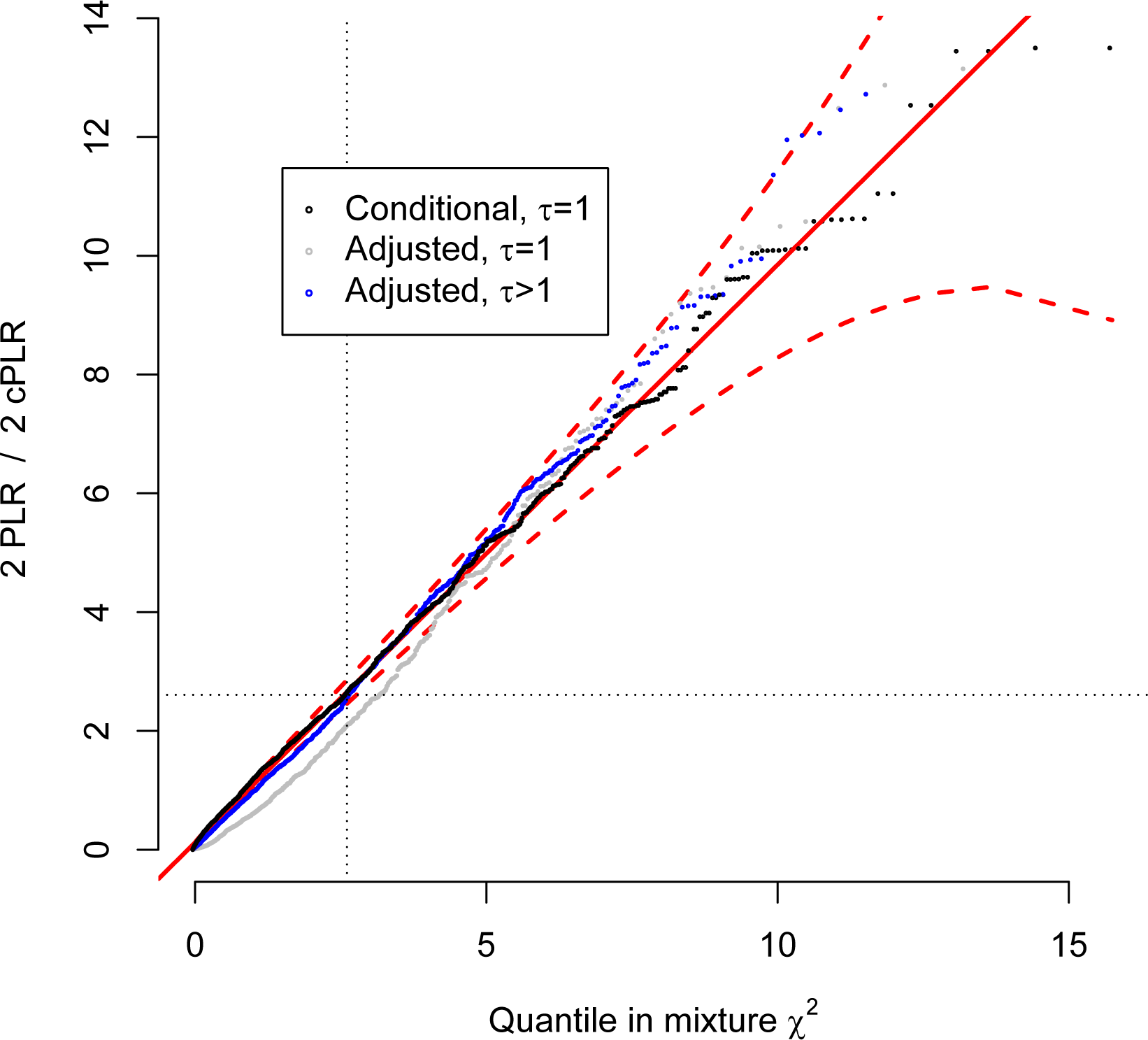
QQ plot from simulations demonstrating type 1 error rate control of PLR test. PLR values for test subgroups under *H*_0_ with either *τ* = 1 (random subgroups; grey) or *τ* > 1 (genetic difference between subgroups, but independent of main phenotype; blue) with cPLR values for random subgroups (black) and against proposed asymptotic distribution under simulation 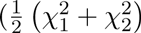; solid red line; 99% confidence limits dashed red line). The distribution of cPLR for random subgroups majorises the distribution of PLR, meaning the PLR-based test is conservative. Further details are shown in supplementary note, section 3.

We then established the suitability of the test when SNPs are in LD and when there exist genetic differences between subgroups that are independent of disease status overall. First, we used a dataset of controls and autoimmune thyroid disease (ATD) cases and repeatedly choose subgroups such that several SNPs had large allelic differences between subgroups. We found good FDR control at all cutoffs (supplementary note, figure 3.2) and the overall type 1 error rate at *α* = 0.05 was 0.041 (95% CI 0.034-0.050). Second, we analysed a dataset of T1D cases with subgroups defined by geographical origin. Within the UK, there is clear genetic diversity associated with region [9]. As expected, *Z_d_* scores for geographic subgroups showed inflation compared to for random subgroups (supplementary figure 1). None of the derived test statistics reached significance at a Bonferroni-corrected *p* < 0.05 threshold (min. corrected p value > 0.8, supplementary figure 2).

To examine the power of our method, we used published GWAS data from the Wellcome Trust Case Control Consortium [10] comprising 1994 cases of Type 1 diabetes (T1D), 1903 cases of rheumatoid arthritis (RA), 1922 cases of type 2 diabetes (T2D) and 2953 common controls. We established that our test could differentiate between any pair of diseases, considered as subgroups of a general disease case group (all < 1 × 10^−8^, table 2).

T1D and RA have overlap in genetic basis [10, 11, 7], as well as non-overlapping associated regions. T1D and T2D have less overlap [11] and T2D and RA less still. This was reflected in the fitted values (table 2, figure 3). The fitted values parametrizing category 2 in the full model for T1D/RA (*π*_2_, *σ*_2_) were consistent with a subset of SNPs associated with case/control status (T1D+RA vs control) but not differentiating T1D/RA. By contrast, the parametrization of category 2 for T1D/T2D and T2D/RA had marginal variance *σ*_2_ approximately 1, suggesting that a subset of SNPs associated with case/control status but not with ‘subgroup’ status did not exist in these cases. The rejection of *H*_0_ for the comparisons entails the existence of a set of SNPs associated both with case/control and subgroup status. The *H*_0_ model does not allow such a set of SNPs, forcing the parametrisation of *Z_d_*, *Z_a_* scores for such SNPs to be ‘squashed’ into a category shape permitted under *H*_0_, with one marginal variance being 1: either category 2 (as happens in T2D/RA since *π*_2_|*H*_0_ ≈ *π*_3_|*H*_1_, *σ*_2_|*H*_0_ ≈ *σ*_3_|*H*_1_ in T2D/RA) or category 3 (as in T1D/T2D, where *π*_3_|*H*_0_ ≈ *π*_3_|*H*_1_, *τ*|*H*_0_ ≈ *τ*|*H*_1_).

**Figure 3:**
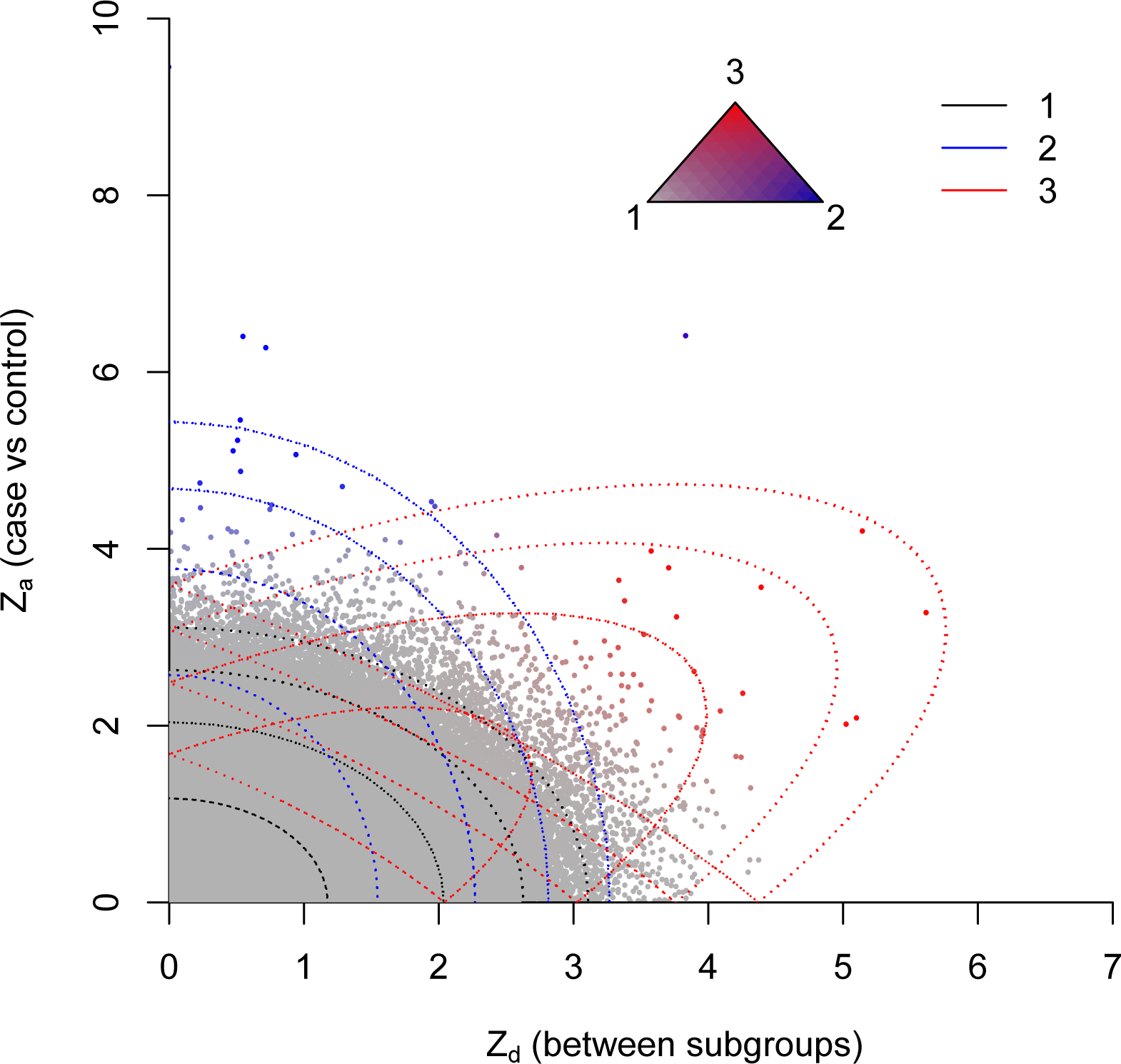
Observed absolute *Z*_*a*_ and *Z*_*d*_ for T1D/RA. Colourings correspond to posterior probability of category membership under full model (see triangle): grey-category 1, blue-category 2, red-category 3. Contours of the component Gaussians in the fitted full model are shown by dotted lines.

**Table 2:** Fitted parameter values for models of T1D/RA, T1D/T2D, T2D/RA, and GD/HT. *H*_1_ is the null hypothesis (under which *σ*_3_ = 1, *ρ* = 0) that SNPs differentiating the subgroups are not associated with the overall phenotype; *H*_1_ is the alternative (full model). *p* values for pseudo-likelihood ratio tests are also shown.

| | | $\pi_1$ | $\pi_2$ | $\pi_3$ | $\sigma_2$ | $\sigma_3$ | $\tau$ | $\rho$ | p-val |
| --- | --- | --- | --- | --- | --- | --- | --- | --- | --- |
| T1D/RA | $H_1$ | 0.997 | $5.69 \times 10^{-4}$ | $2.06 \times 10^{-3}$ | 2.76 | 1.39 | 1.74 | 1.815 | $3.2 \times 10^{-12}$ |
| | $H_0$ | 0.997 | $6.26 \times 10^{-4}$ | $2.48 \times 10^{-3}$ | 2.71 | - | 1.67 | - | |
| T1D/T2D | $H_1$ | 0.573 | 0.426 | $9.63 \times 10^{-4}$ | 1.00 | 2.03 | 2.25 | 1.68 | $1.6 \times 10^{-9}$ |
| | $H_0$ | 0.578 | 0.421 | $8.91 \times 10^{-4}$ | 1.00 | - | 2.21 | - | |
| T2D/RA | $H_1$ | 0.573 | 0.426 | $8.71 \times 10^{-4}$ | 1.00 | 2.23 | 1.75 | 1.69 | $5.1 \times 10^{-9}$ |
| | $H_0$ | 0.91 | $8.05 \times 10^{-4}$ | 0.0892 | 2.25 | - | 0.97 | - | |
| GD/HT | $H_1$ | 0.506 | 0.487 | 0.007 | 1.12 | 2.90 | 1.65 | 2.61 | $2.2 \times 10^{-15}$ |
| | $H_0$ | 0.493 | 0.079 | 0.428 | 1.68 | - | 1.03 | - | |

To determine the power of our test more generally, we showed that power depends on the number of SNPs in category 3 and on the underlying parameters of the true model, depending on the number of samples through the fitted model parameters (Supplementary Note 3.3). We therefore estimated the power of the test for varying numbers of SNPs in category 3 and for varying values of the parameters *σ*_3_, *τ*, and *ρ*. (Figure 4; Supplementary Figure 3). As expected, power increases with an increasing number of SNPs in category 3, reflecting the proportion of SNPs which differentiate case subgroups and are associated with the phenotype as a whole. Power also increases with increasing *τ*, *σ*_3_, and absolute correlation (*ρ*/(*σ*_3_*τ*)) as high values enable better distinction of SNPs in the second and third categories.

**Figure 4:**
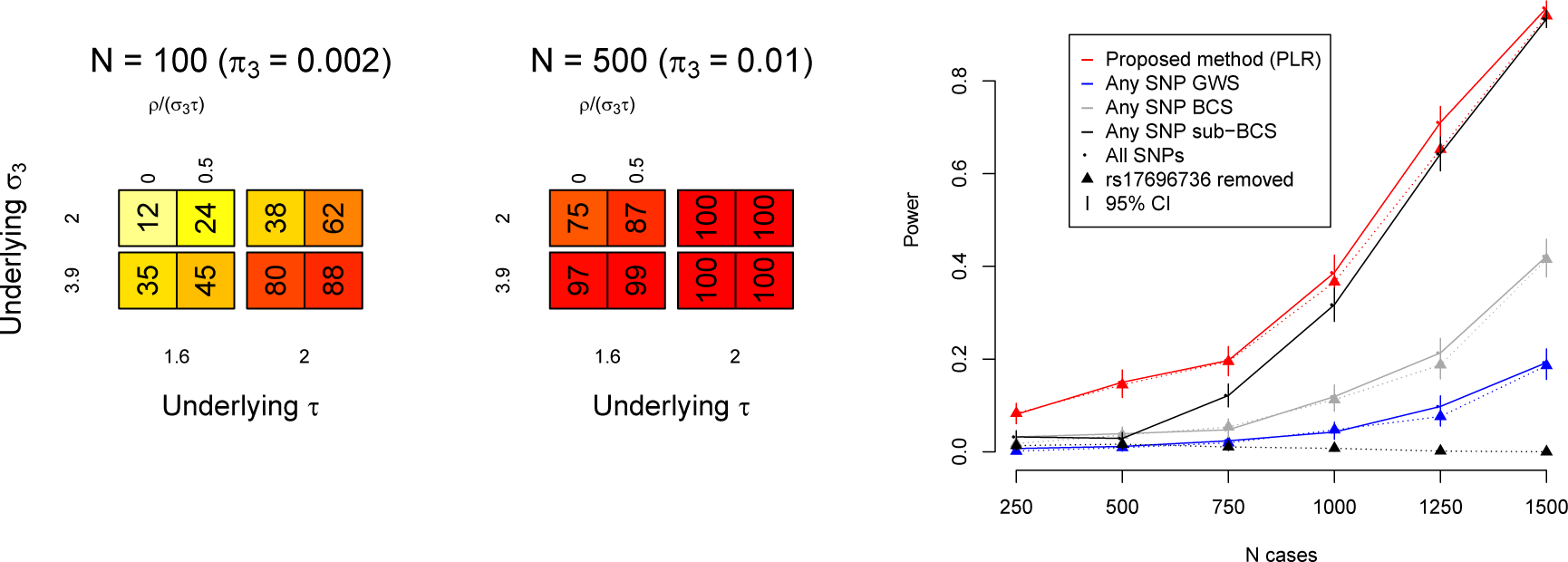
Power of PLR to reject *H*_0_ (genetic homogeneity between subgroups) depends on the number of SNPs in category 3 and the underlying values of model parameters *σ*_2_, *σ*_3_, *τ*, *ρ*. Dependence on number of case/control samples arises through the magnitudes of *σ*_3_ and *τ* (supplementary note, section 3.3). Leftmost figure shows power estimates for various values of *π*_3_, *σ*_3_, *τ*, *ρ*. Value N is the approximate number of SNPs in category 3, (*∝ π*_3_). Each simulation was on 5 × 10_4_ simulated autosomal SNPs in linkage equilibrium. Value *ρ*/(*σ*_3_ *τ*) is the absolute correlation between *Z*_*d*_ and *Z*_*a*_ in category 3. Also see supplementary figure 3. Rightmost figure shows power of PLR to detect differences in genetic basis of T1D and RA subgroups of a combined autoimmune dataset, downsampling to varying numbers of cases (X axis). PLR is compared with: power to find ≥ 1 SNP with *Z*_*d*_ score reaching genome-wide significance (GWS, blue; *p* ≤ 5 × 10^−8^) or Bonferroni-corrected significance (BCS, green; *p* ≤ 0.05?(total # of SNPs)); and power to detect any SNP with *Z*_*a*_ score reaching genome-wide significance and Zd score reaching Bonferroni-corrected significance (sub-BCS, grey; *p* ≥ 0.05?(total # of SNPs with *Z*_*a*_ reaching GWS)). Error bars show 95% CIs. Circles/solid lines for each colour show power for all SNPs, triangles/dashed lines for all SNPs except rs17696736. Power for sub-BCS drops dramatically but power for PLR is not markedly affected, indicating relative robustness of PLR to single-SNP effects.

We explored the dependence of power on sample size by sub-sampling the WTCCC data for RA and T1D (figure 4) and compared the power of the PLR with the power to find any single SNP which differentiated the two diseases in several ways (see figure legend). Although the power of the PLR-based test was limited at reduced sample sizes, it remained consistently higher than the power to detect any single SNP which differentiated the two diseases. We then repeated the analysis removing the known T1D‐ and RA‐ associated SNP rs17696736. The power to detect a SNP with significant *Z_d_* score (Bonferroni-corrected) amongst SNPs with GW-significant *Z_a_* score dropped dramatically, though the power of PLR was only slightly reduced. This illustrated the robustness of the PLR test to inclusion or removal of single SNPs with large effect sizes, a property not shared by single-SNP approaches.

Estimating power requires an estimate of the underlying values of several parameters: the expected total number of SNPs in the pruned dataset with different population MAF in case subgroups, and the distribution of odds-ratios such SNPs between subgroups and between cases/controls. With sparse genome-wide cover, such as that in the WTCCC study, *>* 1250 cases per subgroup are necessary for 90% power (discounting MHC region). If SNPs with greater coverage for the disease of interest are used (such as the ImmunoChip for autoimmune diseases) values of *π*_3_, *σ*_3_ and *τ* are correspondingly higher, and around 500-700 cases per subgroup may be sufficient.

### Application to autoimmune thyroid disease and type 1 diabetes

Autoimmune thyroid disease (ATD) takes two major forms: Graves’ disease (GD; hyperthyroidism) and Hashimoto’s Thyroiditis (HT; hypothyroidism). Differential genetics of these conditions have been investigated. Detection of individual variants with different effect sizes in GD and HT is limited by sample size (particularly HT); however, the *T SH R* region shows evidence of differential effect [12]. T1D is relatively clinically homogenous with no major recognised subtypes, although heterogeneity arises between patients in levels of disease-associated autoantibodies, and disease course differs with age at diagnosis [3]. We analysed both of these diseases.

For ATD, we were able to confidently detect evidence for differential genetic bases for GD and HT (*p* = 2.2×10^−15^). Fitted values are shown in table 2. The distribution of cPLR statistics from random subgroups agreed well with the proposed mixture *χ*^2^ (supplementary figure 4b).

For T1D, we considered four subgroupings defined by plasma levels of the T1D-associated autoantibodies thyroid peroxidase antibody (TPO-Ab, n=5780), insulinoma-associated antigen 2 antibody (IA2-Ab, n=3197), glutamate decarboxylase antibody (GAD-Ab, n=3208) and gastric parietal cell antibodies (PCA-Ab, n=2240). A previous GWAS study on autoantibody positivity in T1D identified only two non-MHC loci at genome-wide significance: 1q23/*FCRL3* with IA2-Ab and 9q34/*ABO* with PCA-Ab [3].

We tested each of the subgroupings retaining and excluding the MHC region. Fitted values for models with and without MHC are shown in supplementary table 2, and plots of *Z_a_* and *Z_d_* scores are shown in supplementary figure 5. Retaining the MHC region, we were able to confidently reject *H*_0_ for subgroupings based on TPO-Ab, IA-2Ab and GAD-Ab (all p-values < 1.0 × 10^−20^). Although there was evidence that SNPs in the dataset were associated with PCA-Ab level (*τ* ≈ 2.5, null model), the improvement in fit in the full model was not significant, and we conclude that such SNPs determining PCA-Ab status are not in general T1D-associated. This can be seen by in the plot of *Z_a_* against *Z_d_* (supplementary figure 5) where SNPs with high *Z_d_* values do not have higher than expected *Z_a_* values.

With MHC removed, the subgrouping on TPO-Ab was significantly better-fit by the full model (*p* = 1.5 × 10^−4^). There was weaker evidence to reject *H*_0_ for GAD-Ab (*p* = 0.002) and IA2-Ab (*p* = 0.008) (Bonferroni-corrected threshold at *α* < 0.05: 0.006). Fitted values of *τ* in both the full and null models for GAD-Ab were ≈ 1, indicating absence of evidence for a category of non-MHC T1D-associated SNPs additionally associated with GAD-Ab positivity. Collectively, this indicates that differential genetic basis for T1D with GAD-Ab and IA2-Ab positivity is driven principally by the MHC region, and although PCA-Ab status is partially genetically determined, the set of causative variants is independent of T1D causative pathways.

The variation in genetic architecture of T1D with age is not fully understood, but previous studies have suggested larger observed effects at known loci in patients diagnosed at a younger age [13, 14, 15, 16]. We investigated whether these differences were indicative of widespread differences in variant effect sizes with age-at-diagnosis, possibly due to differential heritability (see supplementary note 1). We applied the method to T1D dataset with *Z_d_* defined by age at diagnosis (quantitative trait). Fitted values are shown in supplementary table 3 and *Z_a_* and *Z_d_* scores in supplementary figure 6. The hypothesis *H*_0_ could be rejected confidently when retaining or removing the MHC region (p values < 1.0 × 10^−20^ and 0.007 respectively). Signed *Z_d_* and *Z_a_* scores for age at diagnosis showed a visible negative correlation (*p* = 0.002) amongst *Z_d_* and *Z_a_* scores for disease-associated SNPs (*r_g_* method 2, figure 5). This is consistent with a higher genetic liability with lower age at diagnosis.

**Figure 5:**
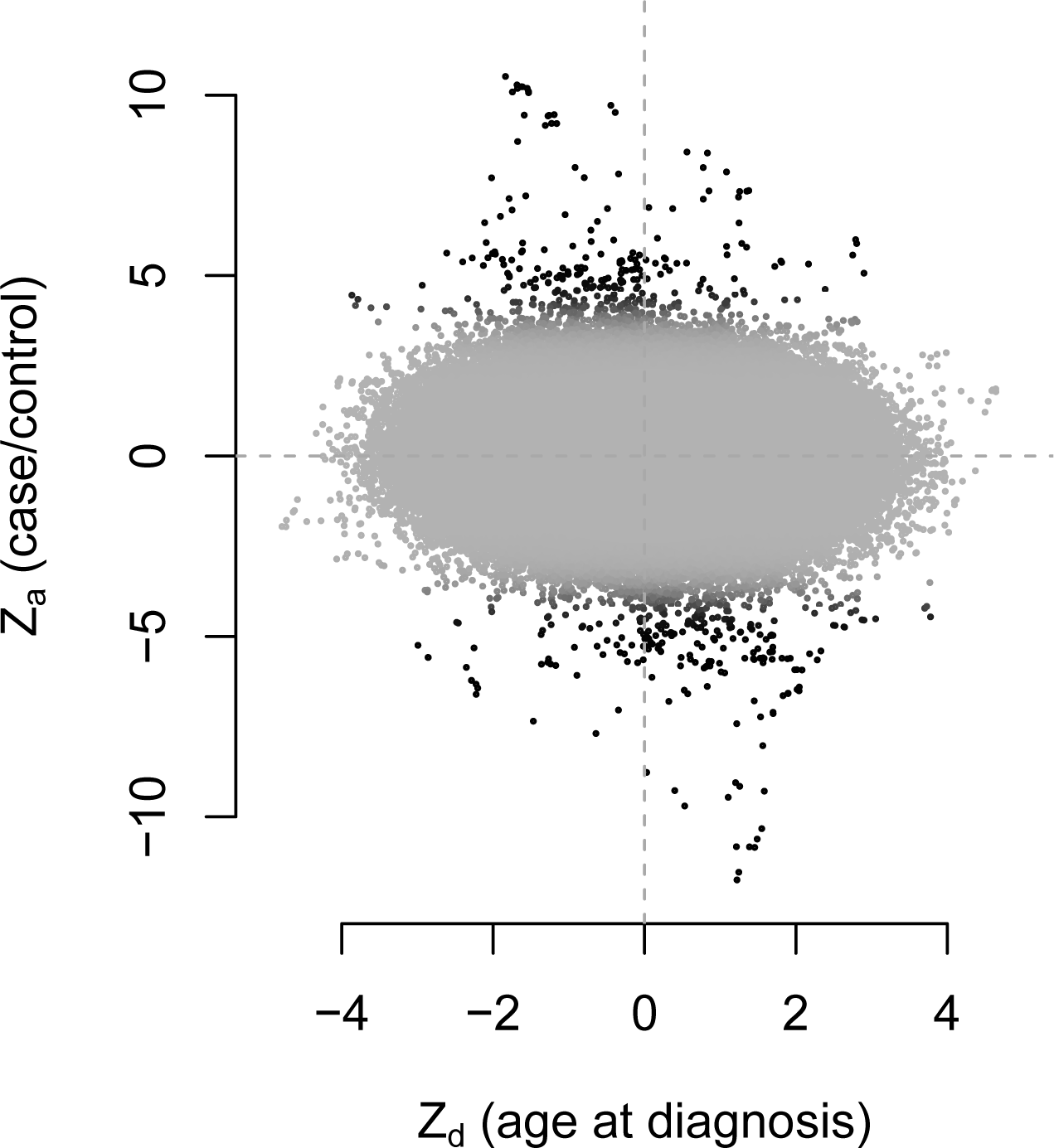
*Z*_*a*_ and *Z*_*d*_ scores for age at diagnosis in T1D, excluding MHC region. Colour corresponds to posterior probability of category 2 membership in null model (since categories in full model are assigned on the basis of correlation), with black representing a high probability. *Z*_*d*_ and *Z*_*a*_ are negatively correlated (*p* = 8.7 × 10^−5^ with MHC included, *p* = 0.002 with MHC removed) after accounting for LD using LDAK weights, and weighting by posterior probability of category 2 membership in the null model, to prioritise SNPs further from the origin

### Assessment of individual SNPs

Many SNPs which discriminated subgroups were in known disease-associated regions (Supplementary Tables 4, 5, and 6). In several cases, our method identified disease-associated SNPs which have reached genome-wide significance in subsequent larger studies but for which the *Z_a_* score in the WTCCC study was not near significance. For example, the SNP rs3811019, in the *PTPN22* region, was identified as likely to discriminate T1D and T2D (*p* = 3.046 × 10^−6^; supplementary table 5), despite a p value of 3 × 10^−4^ for joint T1D/T2D association.

For GD and HT, SNPs near the known ATD-associated loci *PTPN22* (rs7554023), *CTLA4* (rs58716662), and *CEP128* (rs55957493) were identified as likely to be contributing to the difference (see supplementary table 7). The SNPs rs34244025 and rs34775390 are not known to be ATD-associated, but are in known loci for inflammatory bowel disease and ankylosing spondylitis, and our data suggest they may differentiate GD and HT (FDR 0.003).

We searched for non-MHC SNPs with differential effect sizes with TPOA positivity in T1D, the subgrouping of T1D for which we could most confidently reject *H*_0_. Previous work [3] identified several loci potentially associated with TPO-Ab positivity by restricting attention to known T1D loci, enabling use of a larger dataset than was available to us. We list the top ten SNPs for each summary statistic for TPO-Ab positivity in supplementary table 8. Subgroup-differentiating SNPs included several near known T1D loci: *CTLA4* (rs7596727), *BACH2* (rs11755527), *RASGRP1* (rs16967120) and *UBASH3A* (rs2839511) [17]. These loci agreed with those found by Plagnol et al [3], but our analysis used only available genotype data, without external information on confirmed T1D loci. We were not able to replicate the same p-values due to reduced sample numbers.

Finally, we analysed non-MHC SNPs with varying effect sizes with age at diagnosis in T1D (supplementary table 9). This implicated SNPs in or near *CTLA4* (rs2352551), *IL2RA* (rs706781), and *IKZF3* (rs11078927).

## Discussion

The problem we address is part of a wider aim of adapting GWAS to complex disease phenotypes. As the body of GWAS data grows the analysis of between-disease similarity and within-disease heterogeneity has led to substantial insight into shared and distinct disease pathology [6, 7, 2, 20, 21]. We seek in this paper to use genomic data to infer whether such disease subtypes exist. Our problem is related to the question of whether two different diseases share any genetic basis [18] but differs in that the implicit null hypothesis relates to genetic homogeneity between subgroups rather than genetic independence of separate diseases.

Our test strictly assesses whether a set of SNPs have different effect sizes in case subgroups. We interpret this as ‘differential causative pathology’, which encompasses several disease mechanisms, discussed in supplementary note 1. In some cases, if subgroups are defined on the basis of the presence or absence of a known disease risk factor, the heritability of the disease will differ between subgroups, with corresponding changes in variant effect sizes.

We use ‘absolute covariance’ *ρ* preferentially (see supplementary table 1) because we expect that *Z_a_* and *Z_d_* will frequently co-vary positively and negatively at different SNPs in the same analysis; for instance, if some variants are deleterious only for subgroup 1 and others only for subgroup 2. A potential advantage of our symmetric model is the potential to generate *Z_d_* scores from ANOVA-style tests for genetic homogeneity between three or more subgroups, in which case reconstructed *Z* scores would be directionless.

Aetiologically and genetically heterogeneous subgroups within a case group correspond to substructures in the genotype matrix. Information about such substructures is lost in a standard GWAS, which only uses the column-sums (MAFs) of the matrix (linear-order information). Data-driven selection of appropriate case subgroups and corresponding analyses of these subgroups can use more of the remaining quadratic-order information the matrix contains. Indeed a ‘two-dimensional’ GWAS approach (using *Z_a_* and *Z_d_*) instead of a standard GWAS (using only *Z_a_*) may improve SNP discovery, as we found for *PTPN22* in RA/T2D. However, this can only be the case if the subgroups correspond to different variant effect sizes; for other subgroupings, a two-dimensional GWAS will only add noise.

While it seems appealing to use this method to search for some ‘optimal’ partition of patients, we prefer to focus on testing subgroupings derived from independent clinical or phenotypic data. Firstly, it is difficult to characterise subgroupings as ‘better’ or ‘worse’, and no one parameter can parametrise the degree to which two subgroups differ; parameters *π*_3_, *τ*, and *ρ* all contribute, and attempts to test the hypothesis using a single measure such as genetic correlation have serious shortcomings (supplementary note, 4). Secondly, even if subgroups could meaningfully be ranked, the search space of potential subgroupings of a case group is prohibitively large (2*^N^* for *N* cases), making exhaustive searches difficult.

We demonstrated that effect sizes of T1D-causative SNPs differ with age at disease diagnosis. The strong negative correlation observed (figure 5) was consistent with an increased total genetic liability in samples with earlier age of diagnosis, a finding supported by candidate gene studies [14, 15, 16] and epidemiological data [13]. Such a pattern arises naturally from a liability threshold model where total liability depends additively on both genetic effects and environmental influences which accumulate with age (supplementary note 1).

Our method necessarily dichotomises the multitude of mechanisms of heterogeneity, although there are many diverse forms (supplementary table 1, supplementary note 1). There is potential to further dissect the mechanisms of disease heterogeneity by incorporating estimations of genetic correlation [18] or assessing evidence for liability threshold models [22]. Similar mixture-Gaussian approaches may also be adaptable to this purpose, by assessing other families of effect size distributions.

Our method adds to the current body of knowledge by extracting additional information from a disease dataset over a standard GWAS analysis, and determines if further analysis of disease pathogenesis in subgroups is justified. Our approach is analogous to the intuitive method of searching for between-subgroup differences in SNPs with known disease associations [3] but does not restrict attention to strong disease associations, enabling use of information from disease-associated SNPs which do not reach significance. Our parametrisation of effect size distributions allows insight into the structure of the genetic basis of the disease and potential subtypes, improving understanding of genotype-phenotype relationships.

## Methods

### Ethics Statement

This paper re-analyses previously published datasets. All patient data were handled in accordance with the policies and procedures of the participating organisations.

### Joint distribution of variables *Z_a_*, *Z_d_*

We assume that SNPs may be divided into three categories, as described in the results section (figure 1). Under these assumptions, *Z_a_* and *Z_d_* scores have the joint *pdf* given by equation 1. We define Θ is the vector of values (*π*_1_, *π*_2_, *π*_3_, *τ*, *σ*_2_, *σ*_3_, *ρ*). Z scores *Z_a_* and *Z_d_* are reconstructed from GWAS p-values for SNP associations. In practice, since our model is symmetric, we only require absolute Z scores, without considering effect direction.

For sample sizes *n*_1_, *n*_2_ and 97.5% odds-ratio quantile *α*, the expected observed standard deviation of *Z* scores (that is, *σ*_2_, *σ*_3_, and *τ*) is given by

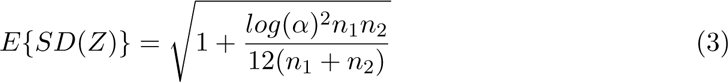

(supplementary note, section 3.3).

### Definition and distribution of PLR statistics

For a set of observed *Z* scores (*Za, Zd*) we define the joint unadjusted pseudo-likelihood *PL_da_*(*Z*|Θ) as

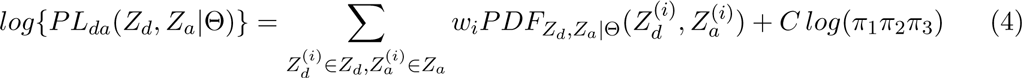

where the term *C log*(*π*_1_*π*_2_*π*_3_) is included to ensure identifiability of the model [5] and weights *wi* are included to adjust for LD (see below).

We now set

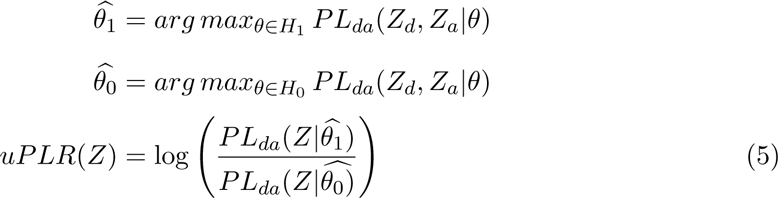

recalling that *H*_0_ is the subspace of the parameter space *H*_1_ satisfying *σ*_3_ = 1 and *ρ* = 0.

If data observations are independent, *uPLR* reduces to a likelihood ratio. Under *H*_0_, the asymptotic distribution of *uPLR* is then

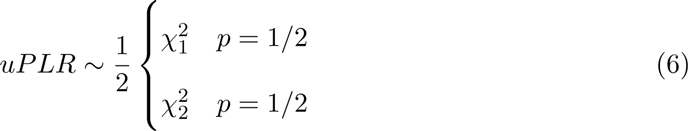

according to Wilk’s theorem extended to the case where the null value of a parameter lies on the boundary of *H*_1_ (since *ρ* = 0 under *H*_0_) [23].

The empirical distribution of *uPLR* may substantially majorise the asymptotic distribution when *τ* ≈ 1. In the full model, the marginal distribution of *Z_a_* has more degrees of freedom (four; *π*_1_, *π*_2_, *σ*_2_, *σ*_3_) than it does under the null model (two; *π*_2_, *σ*_2_; as *σ*_3_ ≡ 1). This can mean that certain distributions of *Z_a_* can drive high values of *uPLR* independent of the values of *Z_d_* (supplementary note 3), which is unwanted as the values *Z_a_* reflect only case/control association and carry no information about case subgroups. If observed *uPLR*s from random subgroups (for which *τ* = 1 by definition) are used to approximate the null *uPLR* distribution, this effect would lead to serious loss of power when *τ* >> 1.

This effect can be managed by subtracting a correcting factor based on the pseudolikelihood of *Z_a_* alone, which reflects the contribution of *Z_a_* values to the uPLR. We define

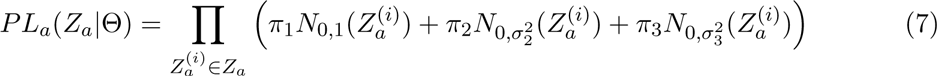

that is, the marginal likelihood of *Z_a_*. Given 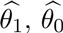 as defined above, we define

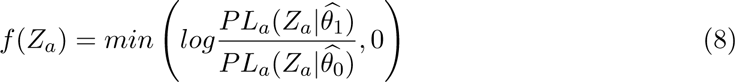

We now define the PLR as

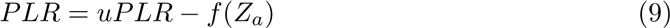

The action of *f*(*Z_a_*) leads to the asymptotic distribution of PLR slightly minorising the asymptotic mixture-*χ*^2^ distribution of uPLR, to differential degrees dependent on the value of *τ* (see supplementary note 3).

We define the similar test statistic *cPLR*:

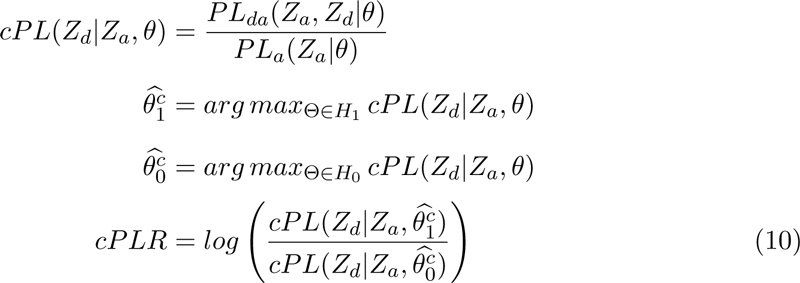

noting that the expression 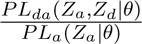 can be considered as a likelihood conditioned on the observed values of *Z_a_*. Now

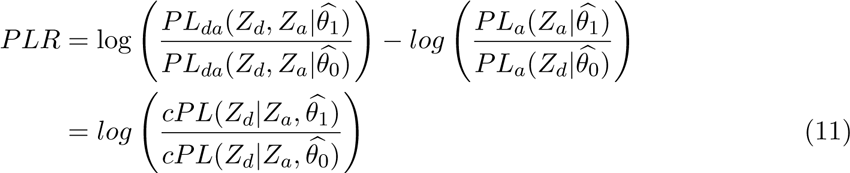

The empirical distribution of cPLR for random subgroups majorises the empirical distribution of PLR (supplementary note 3). Furthermore, the approximation of the empirical distribution of cPLR by its asymptotic distribution is good, across all values of *τ*; that is, across the whole null hypothesis space.

Our approach is to compare the PLR of a test subgroup to the cPLR of random subgroups, which constitutes a slightly conservative test under the null hypothesis (see supplementary note 3).

### Allowance for linkage disequilibrium

The asymptotic approximation of the pseudo likelihood-ratio distribution breaks down when values of *Z_a_*, *Z_d_* are correlated due to LD. One way to overcome this is to ‘prune’ SNPs by hiererarchical clustering until only those with negligible correlation remain. A disadvantage with this approach is that it is difficult to control which SNPs are retained in an unbiased way without risking removal of SNPs which contribute greatly to the difference between subgroups.

We opted to use the LDAK algorithm [4], which assigns weights to SNPs approximately corresponding to their ‘unique’ contribution. Denoting by *ρ_ij_* the correlation between SNPs *i*, *j*, and *d*(*i*, *j*) their chromosomal distance, the weights *w_i_* are computed so that

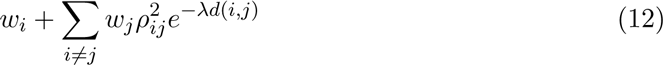

is close to constant for all *i*, and *w_i_* > 0 for all *i*. The motivation for this approach is that, 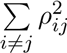 represents the replication of the signal of SNP *i* from all other SNPs.

This approach has the advantage that if *n* SNPs are in perfect LD, and not in LD with any other SNPs, each will be weighted 1*/n*, reducing the overall contribution to the likelihood to that of one SNP. In practice, the linear programming approach results in many SNP weights being 0. Using the LDAK algorithm therefore allows more SNPs to be retained and contribute to the model than would be retained in a pruning approach.

A second advantage of LDAK is that it homogenises the contribution of each genome region to the overall pseudo-likelihood. Many modern microarrays fine-map areas of the genome known or suspected to be associated with traits of interest [24] which could theoretically lead to peaks in the distribution of SNP effect sizes, disrupting the assumption of normality. LD pruning and LDAK both reduce this effect by homogenising the number of tags in each genomic region.

We adapted the pseudo-likelihood function to the weights by multiplying the contribution of each SNP to the log-likelihood by its weight (equation), essentially counting the *i*th SNP *wi* times over. Adjusting using LDAK was effective in enabling the distributions of PLR to be well-approximated by mixture-*χ*^2^ distributions of the form 2 (supplementary plots 4a, 4b, 4c).

### E-M algorithm to estimate model parameters

We use an expectation-maximisation algorithm [25, 26] to fit maximum-PL parameters. Given an initial estimate of parameters 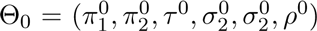 we iterate three main steps:

1. Define for SNP *s* with *Z* scores 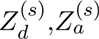

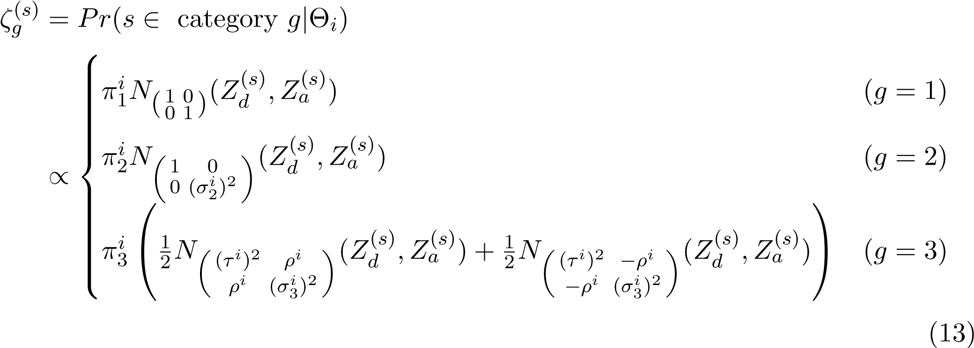
2. For *g* ∈ (1, 2, 3) and LDAK weight *w_s_* for SNP *s* set

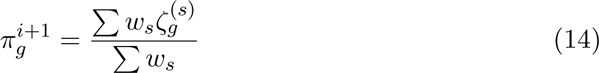
3. Set

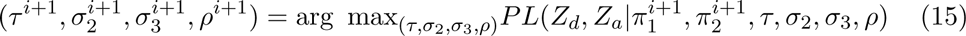

Step 3 is complicated by the lack of closed form expression for the maximum likelihood estimator of *ρ* (because of the symmetric two-Gaussian distribution of category 3), requiring a bisection method for computation. The algorithm is continued until |*PLR*(*Z_d_*, *Z_a_*|Θ_*i*_) − *PLR*(*Z_d_*, *Z_a_*|Θ_*i*−1_)| < ∊; we use ∊ = 1 × 10^−5^.

The algorithm can converge to local rather than global minima of the likelihood. We overcome this by initially computing the pseudo-likelihood of the data at 1000 points throughout the parameter space, retaining the top 100, and dividing these into 5 maximally-separated clusters. The full algorithm is then run on the best (highest-PL) point in each cluster

An appropriate choice of Θ_0_ can speed up the algorithm considerably; for simulations, we begin the model at previous maximum-PL estimates of parameters for earlier simulations.

Maximum-cPL estimations of parameters were made using generic numerical optimisation with the *optim* function in R. Prior to applying the algorithm, parameters *π*_2_ and *σ*_2_ are estimated as maximum-PL estimators of the objective function

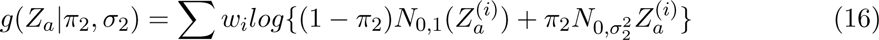

where *w_i_* is the weight for SNP i (see supplementary note 3 for rationale). The conditional pseudo-likelihood was maximised over the remaining parameters.

The algorithm and other processing functions are implemented in an R package available at https://github.com/jamesliley/subtest

### Properties and assumptions of the PLR test

Our assumption that (*Za, Zd*) follows a mixture Gaussian is generally reasonable for complex phenotypes with a large number of associated variants [8] and our adjustment for the distribution of *Z_a_* (essentially conditioning on observed *Z_a_*) reduces reliance on this assumption. If subgroup prevalence is unequal between the study group and population, our method can still be used with adaptation (supplementary note, section 2.4).

Our test is robust to confounders arising from differential sampling to the same extent as conventional GWAS. For example, if subgroups were defined based on population structure, and population structure also varied between the case and control group, SNPs which differed by ancestry would also appear associated with the disease, leading to a loss of control of type-1 error rate. However, the same study design would also lead to identification of spurious association of ancestry-associated SNPs with the phenotype in a conventional GWAS analysis. As for GWAS, this effect can be alleviated by including the confounding trait as a covariate when computing p-values (Supplementary Note 2).

### Prioritisation of single SNPs

An important secondary problem to testing *H*_0_ is the determination of which SNPs are likely to be associated with disease heterogeneity. Ideally, we seek a way to test the association of a SNP with subgroup status (ie, *Z*_*d*_), which gives greater priority to SNPs potentially associated with case/control status (ie, high *Z*_*a*_).

An effective test statistic meeting these requirements is the Bayesian conditional false discovery rate (cFDR) [6]. It tests against the null hypothesis 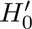 that the population minor allele frequencies of the SNP in both case subgroups are equal (ie, that the SNP does not differentiate subgroups), but responds to association with case/control status in a natural way by relaxing the effective significance threshold on |*Z_d_*|. This relaxation of threshold only occurs if there is systematic evidence that high |*Z_d_*| scores and high |*Z_a_*| scores typically co-occur. The test statistic is direction-independent.

Given a set of observed *Z*_*a*_ and *Z*_*d*_ values 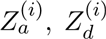, with corresponding two-sided p values *p*_*ai*_, *p*_*di*_, the cFDR for SNP *j* is defined as

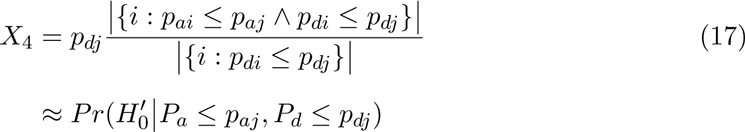

The value gives the false-discovery rate for SNPs whose p-values fall in the region [0, *p_dj_*] × [0, *p_aj_*]; this can be converted into a false-discovery rate amongst all SNPs for whom *X*_4_ passes some threshold [7].

We discuss three other single-SNP test statistics in supplementary note 5.1, which test against different null hypotheses. If the hypothesis 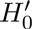 is to be tested, then we consider the cFDR the best of these.

Contour plots of the test statistics for several datasets are shown in supplementary figures 7,8, and 9.

### Genetic correlation testing

Given the correlation between *Z*_*d*_ and *Z*_*a*_ in the age-at-diagnosis analysis, methods to estimate narrow-sense genetic correlation (*r*_*g*_) [18, 19] may be adaptable to the subgrouping question by estimating *r*_*g*_ across a set of SNPs between case/control traits of interest, with the potential advantage of characterising heterogeneity using a single widely-interpretable metric. This may be between *Z* scores derived from comparing the control group to each case subgroup, testing under the null hypothesis *r*_*g*_ = 1 (method 1); or between the familiar *Z*_*a*_ and *Z*_*d*_, under the null hypothesis *r*_*g*_ = 0 (method 2).

We explored these methods in supplementary note 4. We show that method 1 leads to systematically high false positive rates, as *r*_*g*_ is also reduced from 1 in subgroupings that are independent of the overall disease process (e.g. hair colour in T2D). We show that method 2 is considerably less powerful than our method because it tests a narrower definition of *H*_1_ which does not take account of the marginal variances of the distribution of *Z*_*d*_, *Z*_*a*_ in category 3, and requires that correlation between *Z*_*d*_ and *Z*_*a*_ be always positive or always negative, in contrast to our symmetric model (Figure 1). Indeed, parameter *ρ* estimates an analogue of *r*_*g*_ accounting for simultaneous correlation and anticorrelation.

Methods to compute *r*_*g*_ were not explicitly proposed as a method for subgroup testing, and our analysis does not indicate any general shortcomings. However, comparison with *r*_*g*_ based approaches places our method in the context of established methodology, demonstrating the necessity of considering both variance parameters (*τ*, *σ*_3_) and covariance parameters (*ρ*) in testing a subgrouping of interest.

### Description of GWAS datasets

ATD samples were genotyped on the ImmunoChip [24] a custom array targeting putative autoimmune-associated regions. Data were collected for GWAS-like analyses of dense SNP data [12]. The dataset comprised 2282 cases of Graves’ disease, 451 cases of Hashimoto’s thyroiditis, and 9365 controls.

T1D samples were genotyped on either the Illumina 550K or Affymetrix 500K platforms, gathered for a GWAS on T1D [17]. We imputed between platforms in the same way as the original GWAS. The dataset comprised genotypes from 5908 T1D cases and 8825 controls, of which all had measured values of TPO-Ab, 3197 had measured IA2-Ab, 3208 had measured GAD-Ab, and 2240 had measured PCA-Ab. Comparisons for each autoantibody were made between cases positive for that autoantibody, and cases not positive for it. We did not attempt to perform comparisons of individuals positive for different autoantibodies (for instance, TPO-Ab positive vs IA2-Ab positive) because many individuals were positive for both.

To generate summary statistics corresponding to geographic subgroups, we considered the subgroup of cases from each of twelve regions and each pair of regions against all other cases (78 subgroupings in total). To maximise sample sizes, we considered T1D cases as ‘controls’ and split the control group into subgroups.

### Quality control

Particular care had to be taken with quality control, as Z-scores had to be relatively reliable for all SNPs assessed, rather than just those putatively reaching genome-wide significance.. For the T1D/T2D/RA comparison, which we re-used from the WTCCC, a critical part of the original quality control procedure was visual analysis of cluster plots for SNPs reaching significance, and systematic quality control measures based on differential call rates and deviance from Hardy-Weinberg equilibrium (HWE) were correspondingly loose [10]. Given that we were not searching for individual SNPs, this was clearly not appropriate for our method.

We retained the original call rate (CR) and MAF thresholds (MAF ≥ 1%, CR ≥ 95% if MAF ≥ 5%, CR ≥ 99% if MAF <5%) but employed a stricter control on Hardy-Weinberg equilibrium, requiring *p* ≥ 1 × 10^−5^ for deviation from HWE in controls. We also required that deviance from HWE in cases satisfied *p* ≥ 1.91 × 10^−7^, corresponding to |*z*| ≤ 5. The looser threshold for HWE in cases was chosen because deviance from HWE can arise due to true SNP effects [27]. We also required that call rate difference not be significant (*p* ≥ 1 × 10^−5^) between any two groups, included case-case and case-control differences. Geographic data was collected by the WTCCC and consisted of assignment of samples to one of twelve geographic regions (Scotland, Northern, Northwestern, East and West Ridings, North Midlands, Midlands, Wales, Eastern, Southern, Southeastern, and London [10]). In analysing differences between autoimmune diseases, we stratified by geographic location; when assessing subgroups based on geographic location, we did not.

For the ATD and T1D data, we used identical quality control procedures to those employed in the original paper [12, 17]. We applied genomic control [28] to computation of *Z*_*a*_ and *Z*_*d*_ scores except for our analysis of ATD (following the original authors [12]) and our geographic analyses (as discussed above). In all analyses except where otherwise indicated we removed the MHC region with a wide margin (≈ 5*Mb* either side).

## Acknowledgments

We acknowledge the help of the Diabetes and Inflammation Laboratory Data Service for access and quality control procedures on the datasets used in this study. The JDRF/Wellcome Trust Diabetes and Inflammation Laboratory is in receipt of a Wellcome Trust Strategic Award (107212, JAT) and receives funding from the JDRF (5-SRA-2015-130-A-N, JAT) and the NIHR Cambridge Biomedical Research Centre. The research leading to these results has received funding from the European Unions 7th Framework Programme (FP7/2007-2013, JAT) under grant agreement no. 241447 (NAIMIT). JL is funded by the NIHR Cambridge Biomedical Research Centre and is on the Wellcome Trust PhD programme in Mathematical Genomics and Medicine at the University of Cambridge. CW is funded by the Wellcome Trust (089989,107881, CW) and the MRC (MC UP 1302/5, CW). The Cambridge Institute for Medical Research (CIMR) is in receipt of a Wellcome Trust Strategic Award (100140). The funders had no role in study design, data collection and analysis, decision to publish, or preparation of the manuscript.

## Author contributions

AJL: conceived the statistical methods, wrote the software, performed the analyses, analysed the data, and wrote the manuscript. JAT: analysed the data and edited the manuscript. CW: conceived the study, analysed the data, and wrote the manuscript

## Code availability

Code available from https://github.com/jamesliley/subtest (R package)

## Data availability

This paper re-analyses previously published datsets. WTCCC data access for T1D/T2D/RA and controls [10] is described at https://www.wtccc.org.uk/info/access_to_data_samples.html. ATD data are available on request to the original study authors [12]. T1D genetic data from [17] is available at https://www.ncbi.nlm.nih.gov/projects/gap/cgi-bin/study.cgi?study_id=phs000180.v3.p2 which we combined with autoantibody data available from study authors [3]

